# Suppression of MicroRNA-144 Promotes CXCR4 and CXCL12 Expression and Downregulates Apoptosis in Ovarian Cancer Cells

**DOI:** 10.1101/2020.04.17.042382

**Authors:** Fatma Aysun Turut, Hilal Acidereli, Ozge Cevik

**Affiliations:** Department of Biochemistry, Faculty of Pharmacy, Cumhuriyet University, Sivas 58010, Turkey; Department of Biochemistry, School of Medicine, Aydin Adnan Menderes University, Aydin 09100, Turkey

**Keywords:** MiR-144, VEGF, Ovarian cancer, CXCR4, CXCL12, COX-2

## Abstract

MicroRNAs are important regulators in the growth and metastasis of ovarian cancers. Many assays were established to identify the role of miR-144-3p in ovarian cancer cells and its interaction with COX-2 and chemokines (CXCR4 and CXCL12). The ovarian cancer cells (OVCAR-3 and SKOV-3) were transfected with Anti-miR-144 to downregulate the miR-144-3p and cultured for 36 h. We herein examined the cell viability, colony formation, cell migration, COX-2 reporter activity, the protein expressions of CXCR4, CXCL12, COX-2, VEGF, Caspase-3, BAX and Bcl-2. We have observed that the suppression of miR-144-3p significantly increased the cell proliferation and migration and decreased the apoptosis. Moreover, the downregulation of miR-144-3p markedly increased the COX-2, CXCR4, CXCL12 and VEGF expression in OVCAR-3 and SKOV-3 ovarian cancer cells. In conclusion, miR-144-3p may play important roles in the regulation of chemokine receptor CXCR4 and its ligand CXCL12 in the progressive ovarian tumors expressing COX2. These data suggests that miR-144 has the novel therapeutic targets for the cancer therapy and cancer prevention.

## 1. INTRODUCTION

Ovarian cancer is generally a rapidly growing disease and the most lethal of malignant gynecological tumors in women (Lengyel, 2010). Ovarian cancer is a deadly disease, with a cure rate of only 30% in developed countries. One reason for this high mortality rate is the lack of an early detection method for this disease. Ovarian cancer progresses locally in the pelvis and the reproductive organs (uterus, fallopian, tube, ovaries) which are associated with the intraperitoneal organs leading to an early metastasis (Eisenkop and Spirtos, 2001). Therefore, there is a strong need for the prognostic and predictive markers for an early diagnosis and also to optimize a personalized treatment (Buys et al., 2011). Pelvic examination, transvaginal ultrasonography, and the detection of the serum CA125 levels are the standard methods for diagnosing the ovarian tumors. However, they have failed to detect the disease at an early stage and thus reduce the mortality (Myers et al., 2006). As more is discovered about the ovarian cancer biology, our knowledge about the microRNA (miRNA) biology and the expression has grown. miRNAs are predicted to regulate approximately 60% of all human genes providing important clinical, molecular, and genetic knowledge of pathways that contribute to ovarian cancer progression (Kinose et al., 2014). MiRNAs are short non-coded RNAs and involved in posttranscriptional gene regulation. A miRNA can regulate hundreds of mRNAs and multiple miRNAs can regulate an individual mRNA (Lewis et al., 2003).

Up regulation or downregulation of miRNA expression have been found to modulate the expression levels of chemokines and chemokine receptors (Hartmann et al., 2015). The chemokine CXCL12 and chemokine receptor CXCR4 are linked to the proliferation, migration, angiogenesis, metastasis and stem cell mobilization (Liekens et al., 2010). The aim of this study is to evaluate the effect of the suppression of microRNA-144 on the expression of CXCR4 and CXCL12 in ovarian cancer cells and the underlying mechanism about their roles in apoptosis during the metastasis phase of this disease.

## 2. MATERIALS AND METHOD

### 2.1. Cell Culture

OVCAR-3 and SKOV-3 ovarian cancer cells were cultured and maintained in DMEM containing 10% FBS and supplemented with 100 U/mL−1 penicilin and 100 µg/mL streptomycin. The cultures were incubated at 37 °C in a humidified atmosphere with 5% CO2. OVCAR-3 and SKOV-3 cells were seeded in different size plates. Medium was replaced with OptiMEM (Invitrogen, Carlsbad, CA) 24 hours later and the cells were transfected with Anti-miR-144 (hsa-miR-144-3p, AM17100, Invitrogen) (UACAGUAUAGAUGAUGUACU) using Oligofectamine (Invitrogen) at a final concentration of 20 nM. miRNA negative controls (miR-C) were used at the same concentration as the test reagent and the cells were harvested 36 h post-transfection. Celecoxib (CLX), a COX-2 selective inhibitor, was used at COX-2 inhibition (Sigma, PZ0008) and used as positive control.

### 2.2. Cell viability Assay

The effects of Anti-miR-144 and CLX on the viability of ovarian cancer cells were determined using a tetrazolium-based microplate assay with MTT (Vybrant, Life Technologies) (Coruh et al., 2018). Briefly, The OVCAR-3 and SKOV-3 cells were seeded onto a 96-well plate at a density of 1×104 cells/well. Cells were incubated with Anti-miR-144 (0.1-100 nM) and CLX (0.1-100 μM) for 24 h, 48 h and 72 h at different doses. After the incubation, cell culture medium was removed and changed with the plain medium. 20 μL MTT dye (0.5 mg/mL) was added onto each well and incubated for 4 h at 37°C. After that, solubilization buffer (100 μL) was added onto each well and incubated for 1 h at room temperature. Cell viability was determined on a microplate reader (Epoch Biotek Co., USA) at the wavelength of 490 nm and the percentage of living cells were calculated using the Graphpad Prism programme.

### 2.3. Cell growing assays

In order to investigate the effects of Anti-miR-144 on cell survival of OVCAR-3 and SKOV-3 cells, we incubated the cells with Anti-miR-144 at a final concentration of 20 nM in growth medium (without antibiotics) for 36 h. After the incubation, methylene blue staining was used for the determination of the living cell. The cells were extracted with 1% SDS in PBS solution and stained with 0.01% methylene blue solutions. The absorbance was measured at 600 nm with microplate reader. Each experiment was repeated three times.

### 2.4. Determination of COX-2 promoter activity

Cells were seeded onto 24-well plates and co-transfected with totally 0.5 ug plasmids of pCOX-2-Luc (Manvar et al., 2015) and pRL-SV40 (Promega) using the Lipofectamine transfection reagents (Invitrogen). 24 h after transfection, cells were treated with Anti-miR-144 (20 nM) or CLX (20 uM) for another 24 h. At the end of the incubation, cells were lysed in the Passive Lysis Buffer in order to perform the dual luciferase-renilla assays. Luciferase reporter activity was performed on a Luminometer (Lumate, USA) with dual luciferase activity kit (Promega) according to the manufacturer’s protocol.

### 2.5. Colony formation assay

OVCAR-3 and SKOV-3 cells were seeded onto 6-well plate at a concentration of 1000 cells per well, incubated with Anti-miR-144 and maintained at 37°C for 14 days (Cevik et al., 2018). The medium was changed every 3 days. At the end of the incubation period, each well on the plate was washed with PBS, fixed with cold methanol/acetic acid, stained with % 0.5 crystal violet staining solution for 15 min and washed with ddH2O, sequentially. The stained cells were examined with the microscope and the imaging system. The number of colonies in each well was counted and analyzed.

### 2.6. Annexin-V/7AAD staining and flow cytometry

Apoptosis assays were conducted using a flow cytometry-based approach. 2 x105 OVCAR-3 and SKOV-3 cells were seeded on 6 well plates (Matos et al., 2019). The day after seeding, cells were treated with Anti-miR-144 (20 nM) for 36 h. “Annexin-V/Dead Cell kit” (MCH100105, Millipore) was used for detecting cells undergoing apoptosis. Cells were harvested by trypsinisation and resuspended in culture medium. The cells were incubated with “Annexin-V/Dead Cell Reagent” (RT, 20 min, in dark). The live, dead, early and late apoptotic cells were analyzed using Muse Cell Analyzer (Merck Millipore, USA).

### 2.7. Caspase-3 activity assay

In order to determine the cells levels of caspase-3 activity, cells were lysed with cell lysis buffer and centrifuged for 10 min at 9000 rpm at 4°C after treatment of Anti-miR-144 (20 nM) for 36 h (Erdogan et al., 2019). Using a commercial kit (Calbiochem, USA) and following the manufacturer’s instructions, the supernatant was used for measuring caspase-3 activity as a marker of apoptosis in cells.

### 2.8. Protein expression with western blotting

Anti-miR-144 treated cells were collected and lysed with RIPA lysis buffer (included protease inhibitor cocktail) from OVCAR-3 and SKOV-3 cells. Total cellular proteins were measured by bicinchoninic acid assay for the determination of concentrations. SDS-PAGE (12%) gels were prepared and 30 μg protein (equal amount for each well) was loaded into each well for separation. Gels were transferred onto PVDF membranes and the membranes were blocked with 2.5% BSA. After the blocking, the membranes were incubated with specific primary antibodies (COX-2, VEGF, Cas-3, Bax, Bcl-2 and GAPDH, Santa Cruz) at 4°C overnight. Then, the membranes were washed with TBST (containing 0.1% Tween 20) and incubated with the HRP-conjugated secondary antibodies. After that, the membranes were incubated with ECL substrate for the detection of the bands. For the normalization of each band, GAPDH was used as a housekeeping protein as control. Finally, the expression levels of proteins were analyzed and quantified using ImageJ software (NIH, USA).2.9. Quantitative real-time PCR analysis

Total RNA was isolated with RNA extraction kit (Invitrogen) according to the manufacturer’s instructions, as previously described (Cevik et al., 2013).

A total of 1 ug RNA was reverse transcribed as the template for cDNA synthesis using High Capacity cDNA Reverse Transcription Kit (Applied Biosytem) according to the manufacturer’s instructions. Quantitative real-time PCR was performed for COX-2, CXCR4, CXCL12, VEGF and GAPDH using primers. The primers were purchased ready-to-order from KiCqstart Sigma Aldrich. 100 ng of cDNA was amplified using Sybr Green PCR Master Mix (Applied Biosytem) on the ABI StepOne Plus detection system, programmed for 95°C for 10 min, then 40 cycles of: 95 °C for 15 sec, 60 °C for 1 min. For miRNA-144 analysis, RNA was isolated from tissues or cells using a mirVana miRNA Isolation Kit (Ambion, Carlsbad, CA, USA) in accordance with the manufacturer’s instructions. cDNA synthesis real-time PCR was performed using TaqMan miRNA assays per the manufacturer’s recommendations (Applied Biosystems, Foster City, CA). The amplification results were analyzed using StepOne Software v2.3 (Applied Biosystems, Foster City, CA) and the genes of interest were normalized to the corresponding GAPDH results. Data were expressed as fold induction relative to the control.

### 2.10. Statistical analysis

We used the student t-tests or analyses between groups (ANOVA) for statistical analysis as appropriate. Differences were considered significant p-value of 0.05 or less. All data is represented as mean□±□SD, unless otherwise indicated. All experiments were repeated in triplicate. Data from representative experiments are shown.

## 3. RESULTS

In order to investigate the effect of the downregulation of microRNA-144, SKOV-3 and OVCAR-3 ovarian cancer cells were selected with high metastatic potential.

### 3.1. Inhibition of miR-144 cell growth ovarian cancer cells

First, we analyzed the effects of Anti-miR-144 on human ovarian cancer SKOV-3 and OVCAR-3 cell lines. In Figure 1a, cells were transfected with Anti-miR-144 (0.1-100 nM) or control miRNA for 24h. Low concentrations of Anti-miR-144 were able to increase the cell viability to significant levels in both cell lines after 24h of exposure, as shown in dose course experiments, as detected by MTT assay. The increasing percentage of cell viability was significantly different in all groups (p<0.001) when compared with the negative control. On the other hand, Cells were treated with CLX (0.1-100 μM) or control for 24h. Low concentrations of CLX were able to decrease the rate of cell viability in both cell lines significantly after 24h of exposure, as shown in dose course experiments, as detected by MTT assay (Fig 1b).

**Fig 1.**
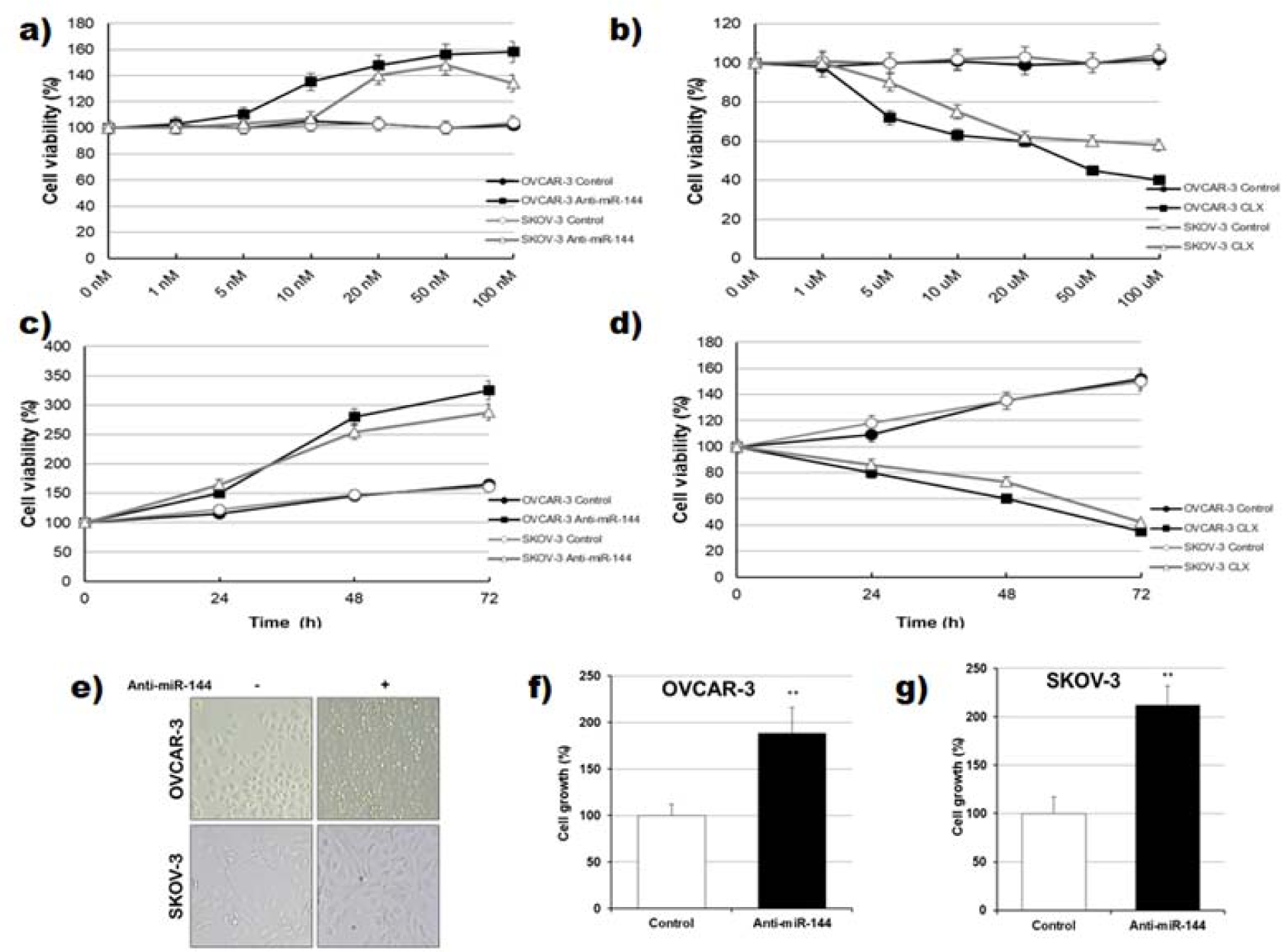
Effects of miR-144 cell survival in ovarian cancer cells. Cell survival measurement with MTT assay after a) Anti-miR-144 treatment (0-100 nM) and b) CLX treatment (0-100 μM) of OVCAR-3 and SKOV-3 cell lines for 36□h. Time dependent effects of c) 20 nM concentration of Anti-miR-144, d) 20 μM concentration of CLX treatment at 24h, 48h, 72h. e) Percentage of cell growing in f) OVCAR-3 and g) SKOV-3 cells with treated of Anti-miR-144 (20 nM) for 36h, (**p□0.01 compare to control)

We next examined whether the cell growth is sustained at different time points (24h, 48h and 72h) with 20 nM Anti miR-144 or 20 μM CLX treatments in SKOV-3 and OVCAR-3 ovarian cancer cells (Fig 1 c-d). Thus, we chose 20 nM Anti-miR-144 or 20 μM CLX with 36 h incubation for further analyzes. As seen in Fig. 1e, SKOV-3 and OVCAR-3 cells treated with 20 nM Anti-miR-144 for 36h showed similar results. The results showed that the cell viability in both SKOV-3 and OVCAR-3 cells were significantly increased with Anti-miR-144 treatment compared to the control groups (p<0.01) (Fig 1 f-g).

### 3.2. Anti-apoptotic effects of Anti-miR-144 on ovarian cancer cells

As indicated in Figure 2a, to detect whether Anti-miR-144 suppresses apoptosis in OVCAR-3 and SKOV-3 cells, we performed western blotting experiment for protein expression level of CAS-3, BAX and BCL-2. The protein expressions of CAS-3 were significantly decreased by the transfection of Anti-miR-144 compared to the control group in SKOV-3 cells (p<0.05, Fig 2b) but were not significantly changed in OVCAR-3 cells. Levels of BAX protein expression were decreased >3-fold in OVCAR-3 and >3-fold in SKOV-3 with Anti-miR-144 compared to control groups (p<0.001, Fig 2c). Levels of BCL-2 protein expression were increased >3-fold in OVCAR-3 with Anti-miR-144 compared to control groups (p<0.001, Fig 2d), but were not significantly changed in SKOV-3 cells.

**Fig 2.**
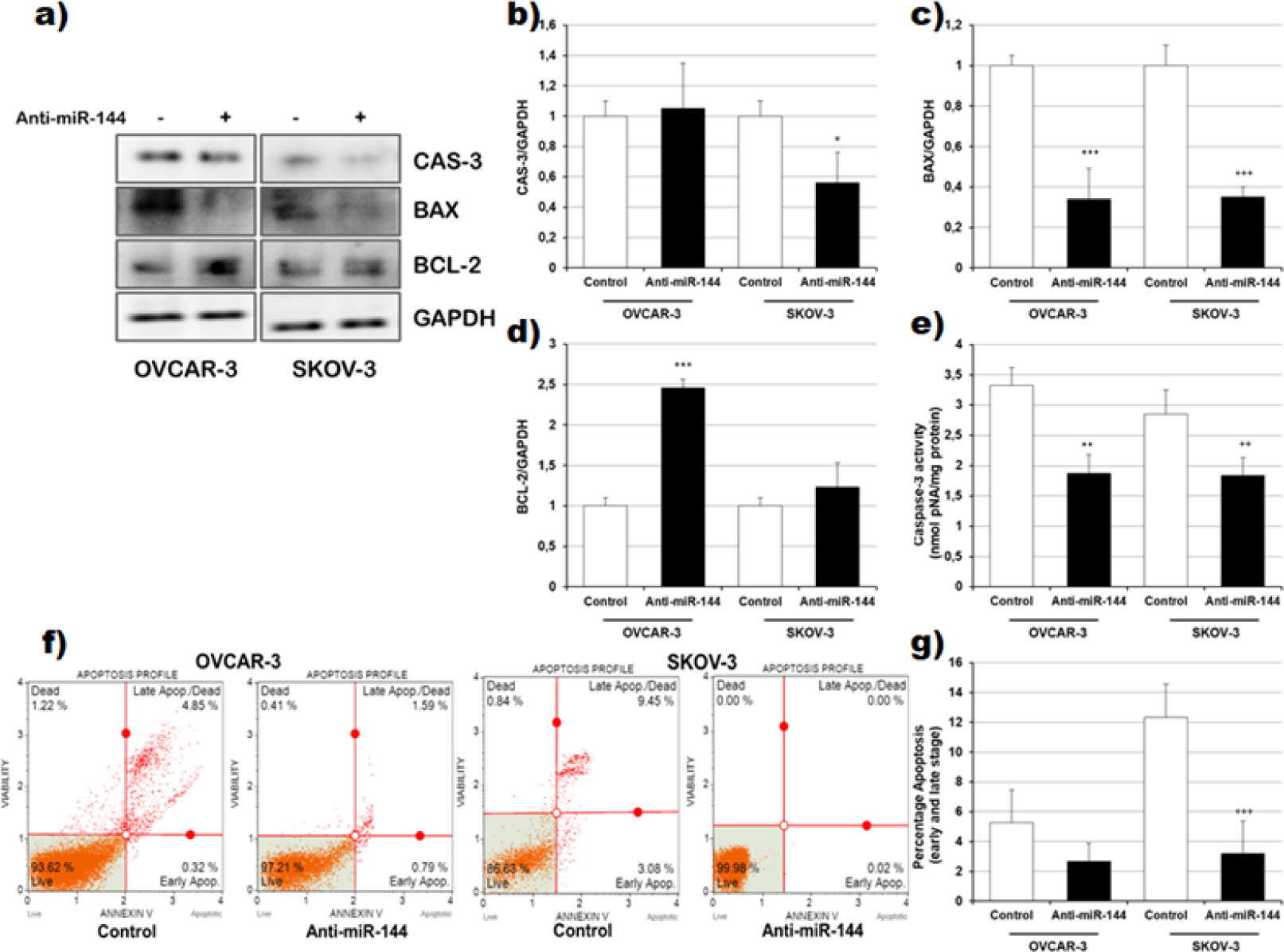
Effects of Anti-miR-144 on apoptotic proteins in ovarian cancer cells. a) Western blot analysis of expression levels of Caspase-3, Bax, Bcl-2 proteins in Anti-miR-144 (20 nM) treatment at 36h in OVCAR-3 and SKOV-3 cells, representatively. Each protein band was normalized to the intensity of GAPDH used. Graph represents the fold-increase upon Anti-miR-144 treatment compared to control. SD is calculated from the mean of 3 independent experiments. Western blot densitometry analysis of expression levels of b) Cas-3, c) Bax and d) Bcl-2 protein in OVCAR-3 and SKOV-3 cells. e) Caspase-3 activity of Anti-miR-144 (20 nM) treatment at 36h in OVCAR-3 and SKOV-3 cells. f) Annexin-V binding using a Muse cell analyzer (live cells (annexin-V negative (lower right quadrant)), early apoptotic cells (positive for annexin-V (lower right corner)), late apoptotic/dead cells (annexin V positive (upper right quadrant)), and non-apoptotic dead cells (upper left quadrant). g) Percentage of total apoptotic cells (early apoptosis + late apoptotic cells) determined by annexin-V positivity in OVCAR-3 and SKOV-3 cells. (*p□0.05, **p□0.01, ***p□0.001 compare to control).

These data demonstrated that miR-144 regulated apoptosis in ovarian cancer cells by mitochondrial proteins. As an indicator of apoptosis, caspase-3 activity in the OVCAR-3 and SKOV-3 cells was decreased in the Anti-miR-144 treated groups with respect to control group (p<0.001, Fig 2e). Annexin-V is a useful marker based on the detection of translocated phosphatidylserine (PS), a hallmark of apoptosis, and when used in combination with another dye, such as the nucleic acid stain 7AAD, mixed populations of apoptotic and non-apoptotic cells can be accurately assessed. When treated with Anti-miR-144 for 36 h, OVCAR-3 and SKOV3 cells exhibited a slight decrease in PS externalization as determined by flow cytometry analysis using Annexin-V/7AAD staining (Fig 2f-g).

### 3.3. Colony forming ability and VEGF expression with Anti-miR-144 in ovarian cancer cells

Firstly, we examined that the colony forming ability for in vitro tumorigenesis. The ability of OVCAR-3 and SKOV-3 cells to generate clones and self-renew was evaluated in a serum-starved culture with transfected of Anti-miR-144 (Fig 3a). In Fig. 3b-c, Anti-miR-144 showed 320-340%, and 250-260% stimulation of clonogenic survival and sphere-forming of tumor cells in OVCAR-3 and SKOV-3 cells (p<0.001, Fig 3b-c).

**Fig 3.**
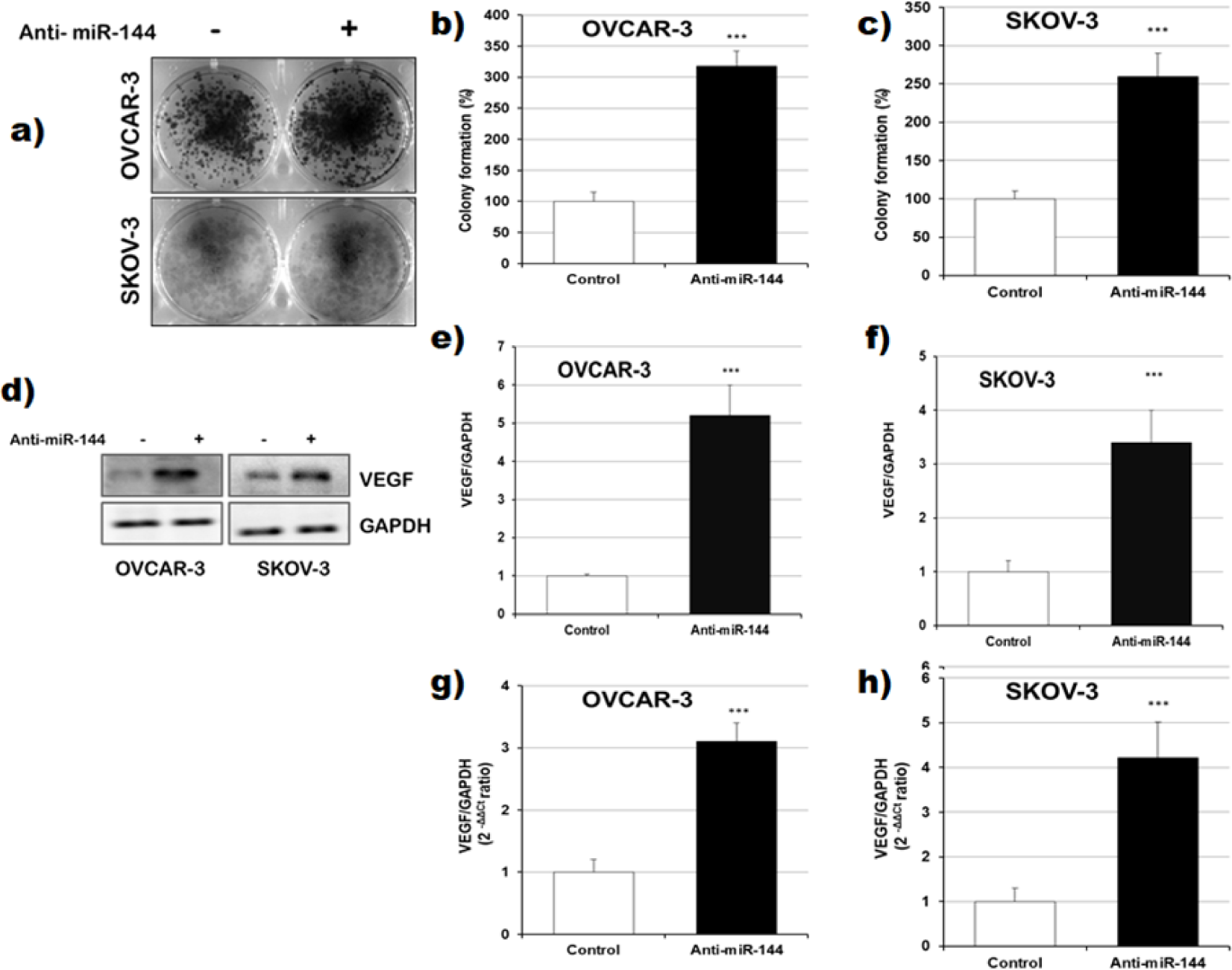
Effects of Anti-miR-144 on cell metastasis in ovarian cancer cells. a) Colony formation assay in OVCAR-3 and SKOV-3 cells. Cells were treated with 20 nM Anti-miR-144 in OVCAR-3 and SKOV-3 cells at 6 well plates and were cultured for 14 days and stained with crystal violet. b-c) Colony formation ratio was quantitated compared to control groups (***p□0.001 compare to control). d) Representative western blot images of VEGF protein expression treated with Anti-miR-144 in OVCAR-3 and SKOV-3 cells. Western blot densitometry analysis of expression levels of VEGF protein in e) OVCAR-3 and f) SKOV-3 cells. (***p□0.001 compare to control). g-h) VEGF gene expression by qPCR analysis of treated with Anti-miR-144 in OVCAR-3 and SKOV-3 cells (***p□0.001 compare to control).

As shown in Fig. 3d, when we exposed OVCAR-3 and SKOV-3 cells to Anti-miR-144 VEGF protein expression was markedly elevated. Levels of VEGF protein expression were increased >5-fold in OVCAR-3 and >3-fold in SKOV-3 cells with Anti-miR-144 compared to control groups (p<0.001, Fig 3e-f). At the same time, levels of VEGF gene expression were upregulated >3-fold in OVCAR-3 and >4-fold in SKOV-3 cells with Anti-miR-144 compared to control groups (p<0.001, Fig 3g-h).

### 3.4. Anti-miR-144 stimulate COX-2 protein expression via COX-2 promoter activity in SKOV-3 and OVCAR-3 cells

We investigated further applications of Anti-miR-144 for COX-2 reporter activity using SKOV-3 and OVCAR-3 cells to study the possible mechanism. The results showed that 20 nM of Anti-miR-144 gradually upregulated and 20 μM CXL, a selective inhibitor for COX-2, downregulated COX-2 promoter activities via luciferase assay, respectively. COX-2 promoter activities with transfection of Anti-miR-144 were increased more than 4 fold in OVCAR-3 cells and and more than 3 fold in SKOV-3 cells, respectively (Fig 4a-b, p<0.001). In order to find out how Anti-miR-144 influenced COX-2 protein and gene expression, we selected 20 nM of Anti-miR-144 transfection for 36 h using western blotting and qPCR. COX-2 gene expression was upregulated in OVCAR-3 (p<0.001) and SKOV-3 (p<0.001) ovarian cancer cells, which were transfected with the Anti-miR-144 (Fig. 4c-d).

**Fig 4.**
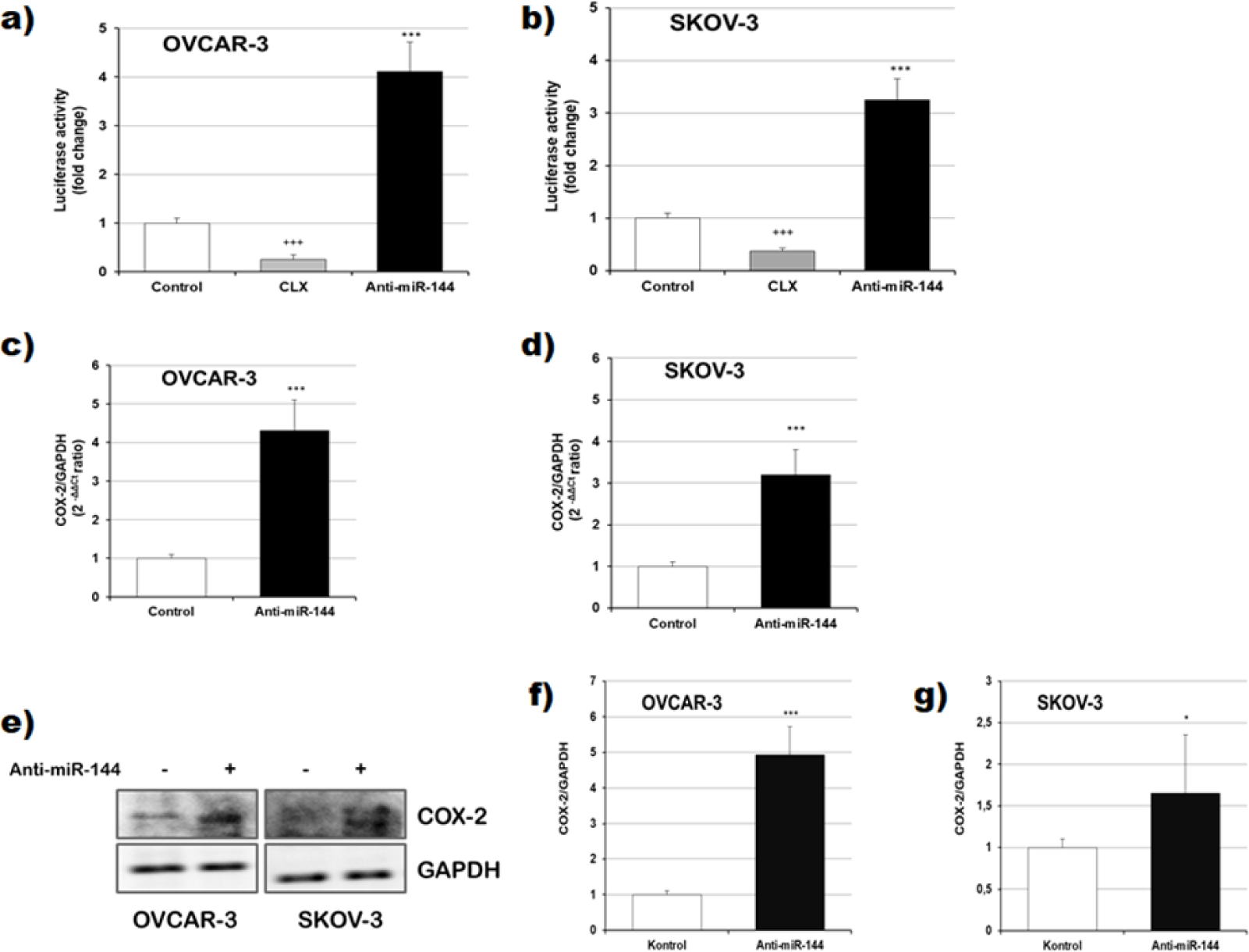
Effects of Anti-miR-144 on COX-2 protein and gene expression in ovarian cancer cells. COX-2 promoter activity in CLX or Anti-miR-144 treated a) OVCAR-3 and b) SKOV-3 cells (***p□0.001, +++p□0.001 compare to control). Reporter firefly luciferase activities were normalized with internal control Renilla luciferase. c-d) COX-2 gene expression by qPCR analysis of treated with Anti-miR-144 in OVCAR-3 and SKOV-3 cells (***p□0.001 compare to control). e) Representative western blot images of COX-2 protein expression for treated with Anti-miR-144 in OVCAR-3 and SKOV-3 cells. Western blot densitometry analysis of expression levels of COX-2 protein in f) OVCAR-3 and g) SKOV-3 cells. (*p□0.05, ***p□0.001 compare to control).

Similarly, there was a remarkable increase in COX-2 protein expression in OVCAR-3 (p<0.001) and SKOV-3 (p<0.001) ovarian cancer cells transfected with the Anti-miR-144 (Fig. 4e-f-g, representative bands were increased more than 2 and 4 fold).

### 3.5. miR-144 and CXCR4/CXCL12 are negatively correlated in ovarian cancer cells

We demonstrated that Anti-miR-144 could upregulate CXCR4/CXCL12 expression. Then we questioned whether overexpression of miR-144 will have an effect on ovarian cancer migration. Firstly, miR-144 gene expression levels were detected with 20 nM Anti-miR-144 transfection or 20 μM CLX incubations in OVCAR-3 and SKOV-3 ovarian cancer cells. Anti-miR-144 downregulated miR-144 gene expression levels in OVCAR-3 (p<0.001, Fig 5a) and SKOV-3 (p<0.001, Fig 5b) ovarian cancer cells. However, CLX gene expression level was upregulated after miR-144 treatment in OVCAR-3 (p<0.001, Fig 5a) and SKOV-3 (p<0.001, Fig 5b) ovarian cancer cells. Next, the gene expression of CXCR4 was significantly elevated with the transfection of Anti-miR-144 and reduced with CLX treatment in OVCAR-3 and SKOV-3 (Fig 5c-d). Moreover, gene expression of CXCL12 was significantly elevated with Anti-miR-144 in OVCAR-3 and SKOV-3 cells (Fig 5c-d). CLX treatment was significantly reduced CXCL12 gene expression in only SKOV-3 cells. (Fig 5e-f). These results suggest that miR-144 is more effective as a chemokine CXCR4/CXCL12 suppressor in cancer cells.

**Fig 5.**
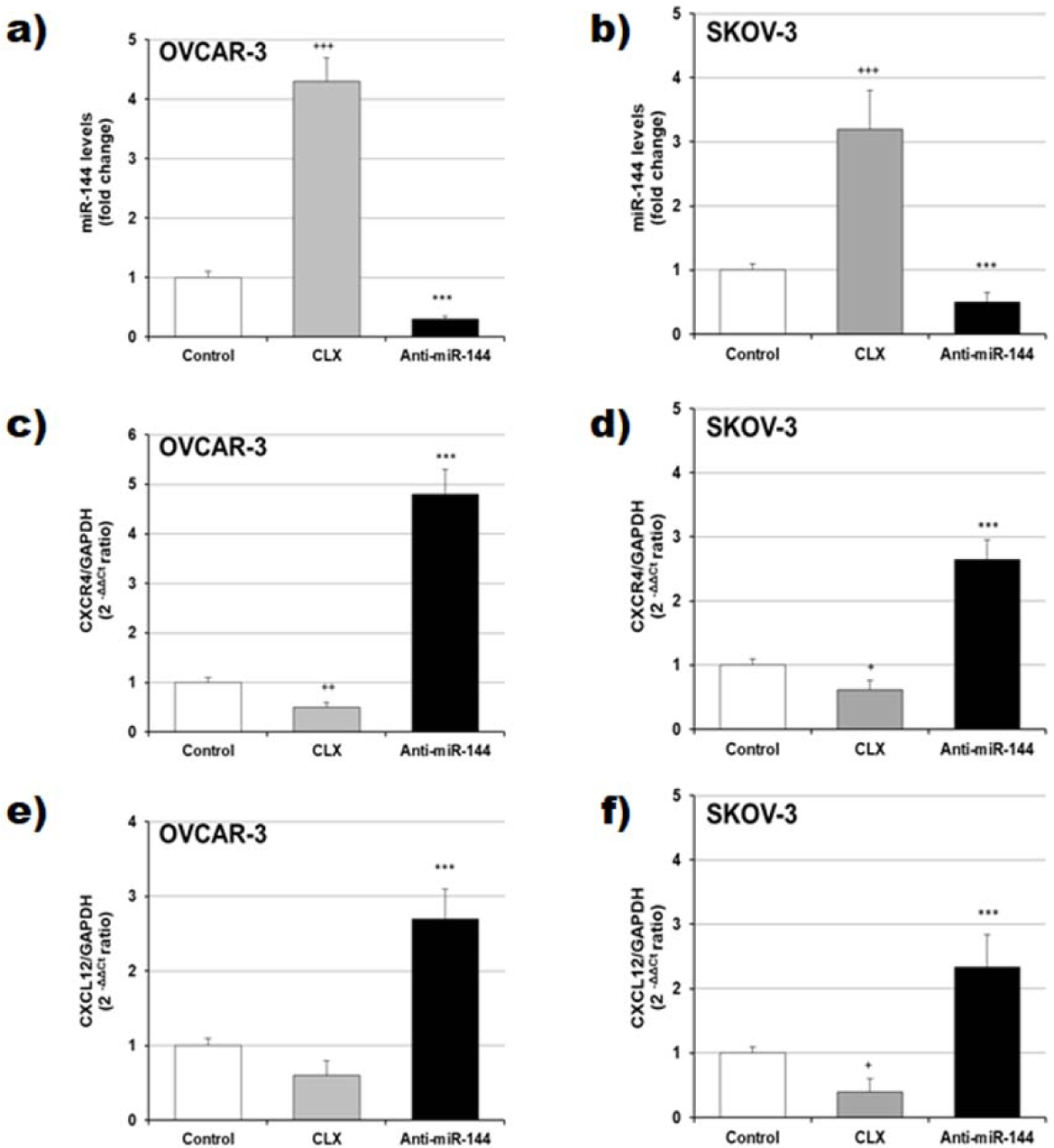
Effects of Anti-miR-144 on CXCR4/CXCL12 cell chemokines in ovarian cancer cells. a-b) miR-144 levels in CLX or Anti-miR-144 treated a) OVCAR-3 and b) SKOV-3 cells (***p□0.001, +++p□0.001 compare to control). c-d) CXCR4 gene expression by qPCR analysis of treated with Anti-miR-144 or CLX in OVCAR-3 and SKOV-3 cells (***p□0.001, +p□0.05 ++p□0.01 compare to control). e-f) CXCL12 gene expression by qPCR analysis of treated with Anti-miR-144 or CLX in OVCAR-3 and SKOV-3 cells (***p□0.001, +p□0.05 ++p□0.01 compare to control).

## 4. DISCUSSION

Ovarian cancer is increasing worldwide and is the leading cause of gynecologic cancer mortality (Webb and Jordan, 2017). Unfortunately, there is no effective early detection to monitor the course of the disease. Most recently, identified serum miRNAs that were correlated with ovarian cancer detection could predict prognosis and chemosensitivity (Vecchione et al., 2013). Several studies analyzed the samples of various ovarian carcinoma and they discovered that different miRNAs were identified to be related to on ovarian cancer (Dahiya and Morin, 2010; Shih et al., 2011). Focusing our attention on miR-144-3p, we unveiled its mechanism of action in modulating chemokines through the metastases on ovarian cancer cells. miR-144 expression was demonstrated to different effects from erythroid lineage to cancer (Dore et al., 2008). At the same time, miR-144 has been observed as a suppressor of cell invasion and metastasis including lymphoma (Wang et al., 2016), hepatocellular (Cao et al., 2014), bladder (Guo et al., 2013) and rectal cancer (Cai et al., 2015). miR-144 is known as a tumor suppressor which inhibits proliferation and promotes apoptosis of osteosarcoma cell lines (Ren et al., 2018). It is recently reported that miR-144-3p can target many oncogenic factors and control autophagy and apoptosis in human lung cancer cells (H292 cells) (Chen et al., 2018). Overexpressing miR-144-3p has inhibited cell proliferation and colony formation, and promoted cell cycle arrest at the G0/G1 phase and apoptosis in multiple myeloma cell lines (Zhao et al., 2017). Moreover, miR-144-3p induced cell cycle arrest and apoptosis in human pancreatic cancer cells (PANC-1 cells) (Li et al., 2017). It is reported that, miR-144 could suppress proliferation, induce cell cycle arrest and cell apoptosis by targeting CEP55 human breast cancer cell lines MCF-7, MDA-MB-231 and SK-BR-3 (Yin et al., 2018). Our results showed that, suppression of miR-144-3p can inhibit mitochondrial apoptosis via downregulation of Bax and upregulation of Bcl-2 and decreasing caspase-3 in human ovarian cancer cells OVCAR-3 and SKOV-3.

Cyclooxygenases (COXs) are cytoplasmic enzymes that catalyze the rate-limiting step of prostaglandin biosynthesis in mediator participating in several biological processes. COX-2 is known to inhibit apoptosis and induce proliferation patterns such as, angiogenesis, inflammation, invasion, and metastasis of cancer cells (Stasinopoulos et al., 2013),(Hashemi Goradel et al., 2019). The present study assessed the suppression of miR-144 is able to markedly activate proliferation and modulate COX-2 expression in ovarian cancer cells. COX-2 as a target for cancer treatment which is known to modulate cell proliferation, cell death, and tumor invasion in many types of cancer. Selective COX-2 inhibitors is used for theraupetic with anti-cancer agents in clinics. Celecoxib is known COX-2-selective inhibitor which use for cancer prevention via anti-tumorigenic effects (Fosslien, 2001b). In this study, we used CLX as a positive control to show COX-2 inhibition. It has been shown that miR-144 down-regulated the synthesis of PGE2 during pregnancy by directly and indirectly inhibiting COX-2 expression in human and mice (Li et al., 2016). In this study, suppression of miR-144 triggered COX-2 protein and gene expression and induced COX-2 promoter activity in ovarian cancer cells. This increase up-regulates the expression of the COX-2 gene, which leads to the increased expression of VEGF (Vascular endothelial cell growth factor). COX-2 and VEGF expressions are highly correlated with increased tumor growth, metastasis and the essential angiogenic proteins of various cancer cells (Fosslien, 2001a). Cancer cells overexpress COX-2 and prostaglandins that reduce cancer cell apoptosis and induce angiogenesis via VEGF (Fosslien, 2001b). Recently, miR-144 acts as a tumor suppressor in the proliferation and metastasis of cervical cancer cells by directly targeting VEGF (Tao et al., 2018). The present study identified with qPCR and western blot analyses that the downregulation of miR-144 may contribute to COX-2 and VEGF expression in ovarian cancers. At the same time, VEGF causes rapid colony formation that causes the proliferation of cancer cells, so when they are VEGF they grow faster than other cancer cells (Yoshida et al., 2015; Lee and Kang, 2018). In this study, when mir-144 was inhibited, capability of a single cell to grow into a large colony of ovarian cancer cells in the cell increased due to COX-2 and VEGF levels.

CXCR4 and its ligand CXCL12 are members of the cXc chemokine family which have roles in tumor proliferation and metastasis in several cancer types (Muller et al., 2001; Kim et al., 2005; Retz et al., 2005). The roles of CXCL12/CXCR4 in ovarian cancer have been discussed in previous studies. Recently, meta-analysis studies showed that CXCL12/CXCR4 expression as a prognostic marker is associated with poor prognosis and influence on survival in ovarian cancer (Liu et al., 2014). CXCR4/CXCL12 pathway and related molecular mechanisms have important roles to play in biological aggressiveness in metastasis and have therapeutic target for epithelial ovarian carcinoma (Jiang et al., 2006; Oda et al., 2007; Sekiya et al., 2012; Li et al., 2014). Certain studies have investigated the role of miRNA expression regulation and interaction with chemokine family as CXCR4/CXCL12 in healthy and cancer cells (Rhodes et al., 2011; Shen et al., 2014; Mei et al., 2017). These studies suggest that CXCR4/CXCL12 signaling stimulated tumorigenesis via microRNA regulation. The present study identified that the suppression of miR-144 may induce the CXCR4/CXCL12 gene expression levels and contribute to metastasis via VEGF expression in ovarian cancer cells. Some authors have shown a significant relationship between VEGF expression and CXCR4/ CXCL12 for metastasis and tumor progression in cancers (Liang et al., 2007; Wang et al., 2011; Miyoshi et al., 2014). On the other hand, CXCL12 and CXCR4 are controlled by the tumor-associated inflammatory mediator prostaglandin E2 (PGE2) in ovarian cancer patients and required COX2 activity. COX2 inhibitors and PGE2 receptor blockers are used to reverse the CXCR4/CXCL12 chemokine responsiveness in the ovarian cancer environment (Obermajer et al., 2011). Another study showed that, COX-2 inhibition suppressed PGE2 production and CXCR4/CXCL12 expression which reduce the formation of stromal tissues around the tumors besides the attenuation of angiogenesis (Katoh et al., 2010). The relationship between CXCR4/CXCL12, COX-2 and VEGF, which has been shown with different studies, shows that it is related to miR-144 in our study.

In conclusion, our results suggested that miR-144-3p suppression may trigger tumor proliferation and angiogenesis and the results revealed that COX-2 and CXCR4/CXCL12 may promote metastasis through the downregulation of miR-144-3p in ovarian cancer. This might shed light on the role of miR-144 in cancer metastasis with chemokine family.

## CONFLICT OF INTEREST

No potential conflicts of interest were identified by any authors of this manuscript.

## ACKNOWLEDGEMENTS

This work was supported by the Cumhuriyet University Research Grant (Projects: ECZ-042) and Science Academy’s Young Scientist Award (BAGEP)-2016 to Dr. Ozge Cevik.

## Notes

### Competing Interest Statement

The authors have declared no competing interest.

